# *SMAP design*: A multiplex PCR amplicon and gRNA design tool to screen for natural and CRISPR-induced genetic variation

**DOI:** 10.1101/2022.07.21.500617

**Authors:** Ward Develtere, Evelien Waegneer, Kevin Debray, Sabine Van Glabeke, Steven Maere, Tom Ruttink, Thomas B. Jacobs

## Abstract

Multiplex amplicon sequencing is a versatile method to identify genetic variation in natural or mutagenized populations through eco-tilling or multiplex CRISPR screens. Such genotyping screens require reliable and specific primer designs, combined with simultaneous gRNA design for CRISPR screens. Unfortunately, current tools are unable to combine multiplex gRNA and primer design into a high-throughput and easy-to-use manner with high design flexibility. Here, we report the development of a bioinformatics tool called *SMAP design* to overcome these limitations. We tested *SMAP design* on several plant and non-plant genomes and obtained designs for more than 80-90% of the target genes, depending on the genome and gene family. We validated the primer designs with Illumina multiplex amplicon sequencing and Sanger sequencing in Arabidopsis and soybean. We also used *SMAP design* to perform eco-tilling by tilling PCR amplicons across nine candidate genes putatively associated with haploid induction in *Cichorium intybus*. We screened 60 accessions of chicory and witloof and identified thirteen knockout haplotypes and their carriers. *SMAP design* is an easy-to-use command-line tool that generates highly specific gRNA and/or primer designs for any number of loci for CRISPR or natural variation screens and is compatible with other SMAP modules for seamless downstream analysis.

## INTRODUCTION

The detection of genetic variation is of great value to medicine and agriculture as it allows researchers to uncover molecular mechanisms, study genetic pathways, and assign gene function. In the frame of breeding, genetic variation can be discovered by screening for beneficial alleles in natural accessions or gene pools. Sequence-based allele mining, also called eco-tilling (1), is a method to screen for naturally-occurring mutations in a set of genomic regions (i.e., candidate genes) across a broad collection of genotypes. A versatile and cost-efficient method for such targeted sequencing is highly multiplex amplicon sequencing (HiPlex), in which multiple target regions (tens to thousands) are amplified in a single PCR reaction and all amplicons are sequenced via Illumina sequencing. Adding sample-specific indices during library preparation allows the pooling of up to hundreds or thousands of samples in a single sequencing run. There are ample examples of studies using this technique in several crops including barley, rice, soy, and wheat (reviewed in Kumar, Sakthivel *et al*. (2)).

Carriers of rare, defective alleles often display useful phenotypes (3, 4) making their identification important for crop breeding and fundamental research. However, population genetics theory predicts that defective alleles may be maintained at a low frequency in natural populations as they negatively affect plant fitness and are subject to negative selection (5). When such carriers are difficult to find in natural populations, genetic variation can be induced with random mutagens like ethyl methanesulfonate or in a targeted fashion with genome editing technologies like CRISPR. CRISPR has become the staple genome editing tool due to its efficacy and simple design. In its most basic form, a CRISPR-associated endonuclease (Cas) is directed to a target site via a guide RNA (gRNA), where it creates a double-stranded DNA break (6, 7). In most eukaryotes, imperfect repair typically results in insertions and/or deletions (indels), or infrequently substitutions, around the breakpoint (8, 9). In principle, CRISPR can be used to knock out any protein-coding gene by disrupting the reading frame or regulatory regions. Multiplex CRISPR screens go a step further by simultaneously targeting multiple genomic loci with arrays of gRNAs to produce large collections of individuals with unique combinations of induced DNA modifications (10). Therefore, eco-tilling and CRISPR are highly complementary mutation screening approaches.

An essential aspect of any mutant screen is the genotyping assay to identify the underlying sequence variant. Specific design parameters need to be considered for each assay’s respective purpose and constraints. For eco-tilling, complete coverage of the regulatory and coding regions is preferred as it allows one to identify all existing mutations in the set of candidate loci across the gene pool and thereby identify conserved and variable genic regions and carriers of defective alleles. While primer design is typically based on a single representative reference genome sequence, eco-tilling may be subject to amplicon dropout in highly divergent regions due to primer-template mismatches. In addition, HiPlex amplicons cannot overlap within a single multiplex reaction as the smaller amplicons, formed through cross-amplification (**Supplementary Figure S1**), would dominate the PCR and reduce coverage. Therefore, in multiplex PCR applications, multiple primer mixtures need to be designed to specifically amplify complementary (partially overlapping) regions across the candidate loci in separate reactions. In contrast, most CRISPR screens do not require complete coverage of the candidate genes. Instead, the gRNA/primer design focuses on specific and efficient gRNAs, using prior knowledge of essential regions of the CDS and/or regulatory sequences, and covers those regions with few amplicons with high primer-binding specificity. Genotyping assays are relatively simple to design manually for a handful of targets, but it can take months for a combinatorial, multiplex CRISPR screen with hundreds of targets. Designing amplicons for gene families is particularly challenging due to sequence similarity between paralogous genes and the chance of cross-amplification or off-target amplification (here collectively called mispriming; **Supplementary Figure S1**). In addition, specific design parameters such as amplicon size range need to be adjusted depending on the downstream library preparation and sequencing technology (e.g., paired-end Illumina short reads or Sanger sequencing). While there are several online and command-line tools available for gRNA design (CRISPOR, CHOPCHOP, FlashFry, CRISPRscan, CCTop) and primer design (PrimerMapper, PrimerView, Primer3), none are integrated with genotyping assay design in a high-throughput manner with the flexibility and specificity required for large-scale multiplex experiments (11–18). Thus, combined gRNA and amplicon design is currently one of the limiting factors to perform medium to high-throughput multiplex CRISPR screens on tens to thousands of genes.

Here, we report the development of a bioinformatics tool called *SMAP design* that addresses these limitations and seamlessly fits into the larger SMAP package that analyzes naturally occurring and CRISPR-induced sequence variants (19). *SMAP design* uses Primer3 (18) to create sets of amplicons with localized or global coverage across reference sequences. For CRISPR experiments, amplicon coordinates are intersected with gRNA target sites from algorithms such as CRISPOR (13) or FlashFry (12), based on user-defined positional boundaries. Sets of amplicons/gRNAs can be created within minutes for tens of loci, or up to a few hours for more complex designs with a few hundred genes. We performed *in silico* designs on 80-95 gene families of varying sizes and in different species and implemented several options to improve success. We further empirically validated the designs by PCR amplification and Sanger sequencing or HiPlex sequencing on reference materials (Arabidopsis and soybean). We performed eco-tilling in natural accessions of chicory and witloof (*Cichorium intybus* var. *sativum* and *C. intybus* var. *foliosum*) using primer sets made by *SMAP design* targeting nine candidate genes putatively involved in haploid induction and demonstrate strategies to enhance the mutation screening capacity by combining multiplexing and sample pooling.

## MATERIAL AND METHODS

### SMAP design

*SMAP design* is a command-line tool written in Python3 and is an addition to the SMAP package (19). The program uses Primer3-py (https://pypi.org/project/primer3-py, version 0.6.1 or newer), Biopython (https://biopython.org, version 1.77 or newer), Pandas (https://pandas.pydata.org, version 1.1.5 or newer), Numpy (https://numpy.org, version 1.18.5 or newer), Matplotlib (https://matplotlib.org, version 3.3.3 or newer). All *SMAP design* runs were performed on a computing cluster with Intel Xeon CPUs on one core. *SMAP design* and its source code are available at https://gitlab.com/ilvo/smap-design, and a detailed user manual and guidelines are available at https://ngs-smap.readthedocs.io/en/latest/design/, under the GNU Affero General Public License v3.0.

### Genome source and selection of candidate genes

All genome and annotation files were retrieved from PLAZA dicot 4.5, PLAZA monocot 4.5, or PLAZA pico 3.0 (**Supplementary Table S1**; (20, 21)). A novel, in-house assembled and annotated reference genome sequence of *C. intybus* var. *sativum* (unpublished) was used for genome-wide identification of gene family members from the selected gene families. Input files for *SMAP design* were generated with *SMAP target-selection. SMAP target-selection* is a command-line tool written in Python3 that extracts the genomic sequence of candidate genes from the reference genome (optionally with extra upstream or downstream flanking regions), based on a user-provided list of gene IDs (optionally combined with grouping based on gene family, genetic network, or genetic pathway membership; here, PLAZA homology groups were used), and orients the sequences in the reference sequence FASTA file so that the CDS is encoded on the positive strand. A GFF file that contains the relative coordinates of the annotated features (gene, CDS, exon, optionally critical domains) of extracted genes is also generated for further downstream analysis with *SMAP design* and the other modules in the SMAP package (19).

### gRNA design with FlashFry and CRISPOR

FlashFry (12) was used to design gRNAs for all the analyses except for the genome-wide design of *Physcomitrium patens*, for which gRNAs were designed by CRISPOR (13). Both programs were run with the default settings except for the mismatch parameter of CRISPOR, which was set to 3 (*--mm*). We used SpCas9 with NGG PAM sequence for all designs.

### SMAP design parameter settings

Primer3 default settings (18) were used with the exception of the user-defined settings in *SMAP design* (**Supplementary Table S2**) and the minimum distance between adjacent forward and reverse primers (set to 5 bp). Four general designs were performed for this report: 1) Design_HiPlex_, the amplicon size was set to 120 – 150 bp and a distance of 15 bp between the gRNAs and primers to be compatible with the HiPlex sequencing service of Floodlight Genomics LLC. 2) Design_PE_, the amplicon size was set to 220 – 250 bp and a 15 bp distance between the gRNAs and primers to be compatible with paired-end 150 bp Illumina sequencing. 3) Design_Sanger_, the amplicon size was set to 400 – 800 bp and a 150 bp distance between the gRNAs and primers as these are the preferred settings for ICE (22) or TIDE (23) analysis of Sanger sequences. 4) Design_NatVar_, the amplicon size was set to 120 – 150 bp and the maximum number of amplicons for each of the candidate genes was requested (**Table 1**). After one design round with Design_NatVar_, the selected primer binding sites were encoded as ‘N’ sequences in the reference sequence input file to exclude those regions from Primer3 design in a second iteration with the same settings. This provides a strategy to iteratively create two or more complementary HiPlex assays with partially overlapping (i.e., tiled) amplicons that together increase coverage of the reference sequence when performed in parallel.

**Table 1:** Primer performance in pooled and individual sequencing of nine chicory genes across accessions of *Cichorium intybus* var. *sativum* and *C. intybus* var. *foliosum*. The *C. intybus* var. *sativum* genotype L9001 was used to create the reference genome sequence and the primer design.

| Dataset | Species | # samples | # amplicons | # amplicons >90% of samples successful | # amplicons <90% and ≥50% of samples successful | # amplicons <50% of samples successful |
| --- | --- | --- | --- | --- | --- | --- |
| Pool | Reference | 6 |  | 87 (99%) | 0 (0%) | 1 (1%) |
|  | var. *sativum* | 104 | 88 | 86 (98%) | 2 (2.3%) | 0 (0%) |
|  | var. *foliosum* | 55 |  | 68 (77%) | 17 (19%) | 3 (3.4%) |
| Individual | Reference | 2 |  | 84 (95%) | 1 (1.6%) | 3 (3.4%) |
|  | var. *sativum* | 156 | 88 | 82 (82%) | 14 (16%) | 1 (1%) |
|  | var. *foliosum* | 140 |  | 65 (73%) | 18 (20%) | 5 (5.7%) |

### Plant material and DNA extraction

DNA was extracted from *Arabidopsis thaliana* Colombia-0 leaves according to Berendzen (24) with the following modifications: extraction buffer was added after the grinding step followed by incubation for 20 minutes at 60 °C and centrifugation for 2 minutes at 1,800 × *g*. DNA was extracted from soybean (*Glycine max*) Williams 82 leaves as previously described (25), with the following modifications: an adapted extraction buffer was used (100 mM Tris HCl (pH 8.0); 500 mM NaCl; 50 mM EDTA; 0.7% (w/v) SDS) and a 70% (v/v) ethanol washing step was included. For chicory (*C. intybus*), tissue pools were created by sampling a leaf punch of each individual in the same Eppendorf tube. Pooled leaf material was ground and homogenized and used for DNA extraction with a CTAB extraction protocol (26). Six leaf punches per individual were used for the individual samples. A detailed description of the chicory plant material, including accession names, can be found in **Supplementary Table S3**.

### DNA sequencing

For Sanger sequencing of Arabidopsis amplicons, PCR was performed with Red Taq DNA Polymerase Master Mix (VWR Life Science) according to the manufacturer’s instructions and purified with magnetic beads (CleanNGS). The PCR amplified regions were Sanger sequenced via Mix2Seq (Eurofins Genomics).

Sets of HiPlex amplicons were designed for Arabidopsis (40 amplicons), soybean (40 amplicons), and chicory (two assays with 45 and 49 amplicons, respectively). Genomic DNA for each species was submitted for HiPlex sequencing (Floodlight Genomics LLC). For Arabidopsis and soybean, the sequencing was done on 24 biological replicates of the reference genotypes. Six technical replicates of the genotype L9001 (*C. intybus* var. *sativum*) were used as controls for the pooled sequencing run and two technical replicates of the L9001 reference genotype were included in the individual plant sequencing run. Details of the pooling strategy can be found in **Supplementary Table S3**.

### Sequence data analysis

Sanger sequence analysis was performed with Geneious Prime 2022.0.1 (https://www.geneious.com). BWA-MEM 0.7.17 (27) was used for HiPlex read mapping with default parameters using the gene targets as the reference sequence. *SMAP haplotype-window* (19) was used for the analysis of the mapped Arabidopsis and soybean HiPlex data with the default parameters except for the minimum read count (*--min_read_count*) which was set to 30 and minimum haplotype frequency which was set to 5 (*- -min_haplotype_frequency*). For the pooled chicory samples, *SMAP haplotype-window* was used with default parameters, except for the minimum read count which was set to 30, minimum haplotype frequency which was set to 4, and the mask frequency which was set to 1 (*--mask_frequency*). The same parameters were used for the individual chicory samples and parameters for discrete calls were set to “dosage” (*--discrete_calls*), dosage filter was set to 2 (*--dosage_filter*) and the frequency interval bounds was set to “diploid” (*--frequency_bounds*). The analysis of the chicory haplotypes revealed that six primer pairs had off-target amplification of pseudogenes and/or genes with similar domains outside the gene family. This issue was resolved by including these (pseudo)genes into the reference FASTA and repeating the read mapping, thereby allowing the reads to map onto their correct reference sequence.

*SMAP effect-prediction* (19) was used to predict the effect of the mutations on the encoded protein of all non-reference haplotypes detected in the HiPlex data of chicory with default parameters. We defined the effect of haplotypes as “mild effect” if more than 50% of the resulting protein sequence was identical to the reference protein and as “strong effect” if less than 50% of the protein sequence was identical to the reference protein.

## RESULTS

### The *SMAP design* workflow

#### Input for SMAP design

*SMAP design* was created to easily and rapidly design sets of multiplex amplicon sequencing primers and gRNAs for small to large-scale CRISPR screens (**Figure 1**). Prior to running *SMAP design*, a FASTA file with sets of candidate gene reference sequences (e.g., entire gene families, gene networks, genetic pathways, or any other customized grouping) can be extracted from the reference genome using *SMAP target-selection. SMAP target-selection* orients all genes with the CDS on the positive strand for a consistent coordinate system and automatically generates a corresponding GFF file with the relative location of gene features (e.g., CDS or critical domains). The FASTA and GFF files are used as input for *SMAP design* and downstream analyses with other modules of the SMAP package (19). If the user wants amplicons to specifically cover one or more gRNAs, a list of gRNAs is provided as a tab-delimited file in the format as described in the user manual (output gRNA files from CRISPOR and FlashFry can be directly fed to *SMAP design*).

**Figure 1:**
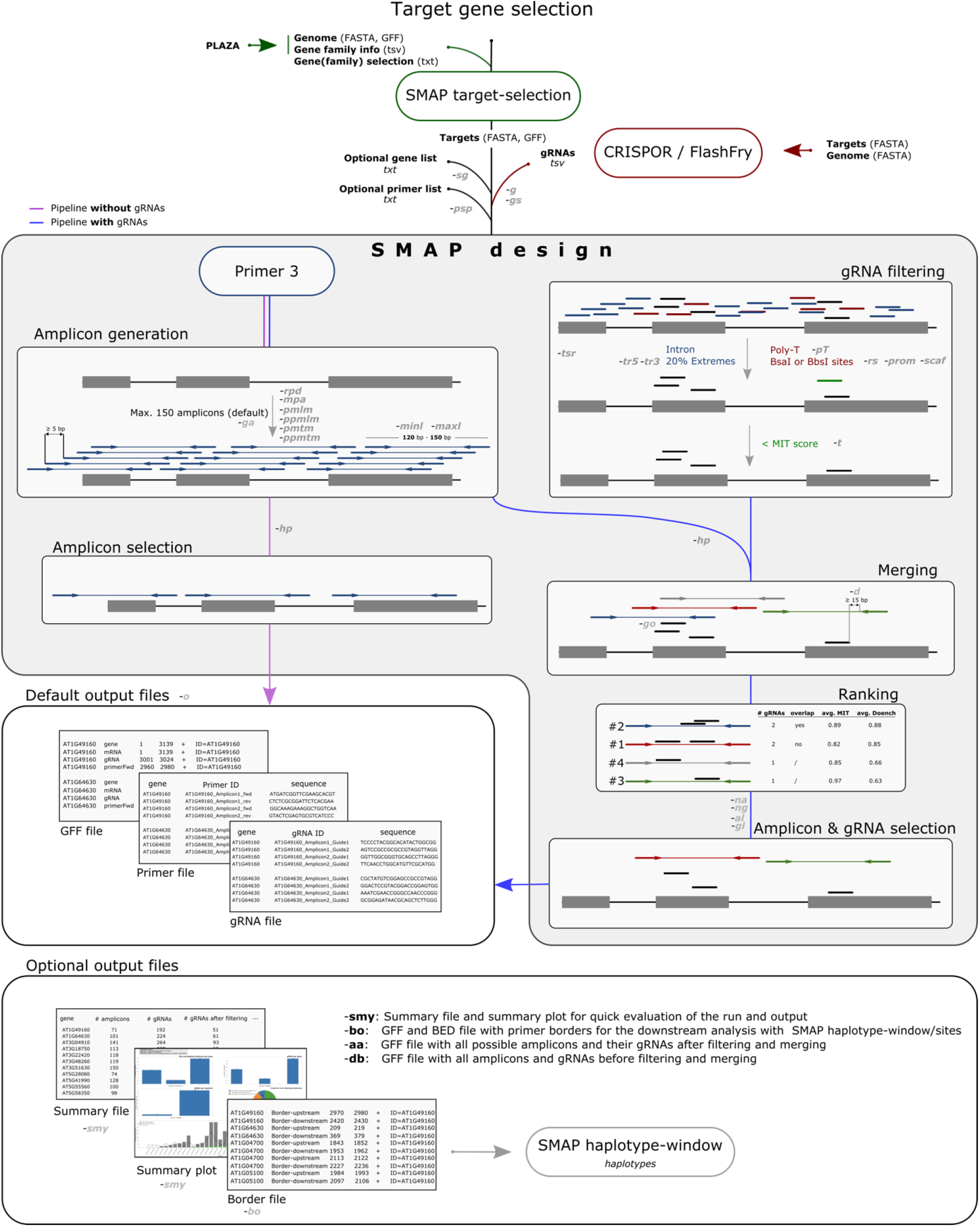
Workflow of *SMAP design*. Users select and extract a set of genes using *SMAP target-selection*. Input files for *SMAP target-selection* can be obtained through PLAZA (or other databases). The FASTA and GFF files are required inputs for *SMAP design*. If no gRNA file is given (purple workflow), *SMAP design* will select only amplicons that do not overlap using amplicons designed by Primer3. If a gRNA file is specified (blue workflow), which can be obtained from third-party software such as CRISPOR or FlashFry, *SMAP design* will filter the gRNAs based on their location in the gene, their sequence, and specificity score. Amplicons designed by Primer3 are merged with the filtered gRNAs and are subsequently ranked based on the gRNAs it overlaps with (number of gRNAs, overlap between gRNAs, and specificity and efficiency scores). Based on the ranking, a maximum (user-defined) number of non-overlapping amplicons per gene are selected. Two or three output files are created by default: a primer and gRNA file with the respective sequences per gene, and a GFF file specifying the location of the primers and gRNAs. Multiple optional files can be generated: a summary file and graph, two debug files, and a border file which is required as input for *SMAP haplotype-window/sites* (for downstream sequence analysis).

#### Amplicon design

*SMAP design* uses the Primer3 module to design, by default, a maximum of 150 amplicons (300 primers) of a user-defined size range for each reference sequence. By default, the specificity of each primer is tested against all reference sequences in the FASTA file to avoid mispriming (**Supplementary Figure S1**). Primer specificity thresholds can be adjusted or switched off. Alternatively, the user can provide a list of gene IDs to limit amplicon design to only that subset, while still using all reference sequences in the FASTA file for primer specificity testing. As Primer3 automatically avoids primer design at ambiguous nucleotides, known polymorphic positions such as SNPs can *a priori* be substituted by N-encoded nucleotides in the reference sequence FASTA to circumvent inefficient primer binding. Primers are spaced by a minimum of 5 bp to spread the amplicons across the target sequences. By default, amplicons with homopolymers (≥ 10 repeated nucleotides) are filtered out because downstream sequencing will likely yield low-quality reads. If no gRNAs are provided by the user (e.g., in case of screening for natural variation), *SMAP design* selects sets of non-overlapping amplicons to maximize reference sequence coverage.

#### gRNA filtering

*SMAP design* filters the provided gRNAs based on several criteria (**Figure 1**). gRNAs with a poly-T stretch (≥4T, a Pol III termination signal) are removed. Short vector sequences directly flanking the gRNA sequence (i.e., promotor and scaffold) can be provided to simulate vector construction steps and exclude gRNAs with restriction sites (e.g., BsaI or BbsI) that interfere with cloning. To increase the likelihood of making knockout mutations, gRNA selection can be focused on selected domains defined via a particular feature type in the annotation GFF (e.g., kinase domain). The user can also exclude a segment of the 5’ and 3’ of the CDS to steer gRNA target sites to a part of the CDS. An optional minimum gRNA specificity score (e.g., MIT score (28)) threshold can also be applied.

#### Ranking and filtering amplicons and gRNAs

Filtered gRNAs are grouped to amplicons by positional overlap. By default, a gRNA is only grouped to an amplicon if the distance between the end of the primer and the gRNA binding site is at least 15 bp. Amplicons are ranked based on the gRNAs they cover according to the following criteria and order: 1) the number of gRNAs (an amplicon with multiple gRNAs will rank higher than an amplicon with a single gRNA); 2) the positional overlap between gRNAs (amplicons with non-overlapping gRNAs will rank highest); 3) the average gRNA specificity scores (e.g., MIT score (28)); and 4) the average gRNA efficiency scores (such as the Doench (29) and out-of-frame scores (30)). If no specificity or efficiency scores are provided in the gRNA file, amplicons are only ranked by the first two criteria. Ultimately, *SMAP design* selects a (user-defined) maximum number of top-ranking, non-overlapping amplicons per gene, each covering a (user-defined) maximum number of gRNAs (**Supplementary Table S2)**.

#### Output of SMAP design

*SMAP design* generates two files by default: a tab-separated values (TSV) file with the primer sequences sequentially numbered per gene and a GFF file with the primer locations on the target gene reference sequences (and other annotation features that were included in the GFF input file). If a gRNA list is provided, *SMAP design* also generates a TSV file with the selected gRNA sequences per gene (**Figure 1**). If no design was possible, the underlying reasons are included per gene at the end of the TSV files. Optionally, summary tables and graphs are generated for a quick evaluation of the set of amplicons and gRNAs (**Supplementary Figure S2**). These graphs show the distributions of the number of gRNAs and non-overlapping amplicons per gene that *SMAP design* generated and indicate the reasons for dropout per gene. For instance, the design may fail because no gRNAs were designed for that gene, none of the gRNAs passed all filters, Primer3 was not capable of designing specific amplicons for the gene, or there was no overlap between the gRNAs and the amplicons. Optionally, a GFF file is created with positions of border sequences required for downstream amplicon analysis by *SMAP haplotype-window* and a BED file required for *SMAP haplotype-sites* (19). In debug mode, an extra GFF output file containing all amplicons and gRNAs prior to filtering is given as a way to visualize the relative positions of all amplicons. Finally after filtering for each gene, an optional GFF file can be generated with all amplicons and their respective gRNAs to visualize and manually select amplicons of interest with a sequence analysis viewer.

### *In silico* testing of *SMAP design* in various species

We evaluated the performance of *SMAP design*, specifically to determine the relationship between successful design and gene family sizes, with the hypothesis that amplicon and/or gRNA designs would be more difficult in larger gene families due to sequence homology. Therefore, we tested *SMAP design* on eleven different species representing a broad range of genome size and compositions (Arabidopsis, *P. patens*, rice, tomato, potato, maize, soybean, Chlamydomonas, *Saccharomyces cerevisiae*, mouse, and human). Per species, 80 - 95 gene families (PLAZA homology groups (20, 21)) (**Supplementary Table S4)**, containing between 1 and 448 genes per family were selected. Three different design settings (Design_HiPlex_, Design_PE_ and Design_Sanger_, see Material and Methods) for different genotyping approaches were tested on the various genes (**Figure 2**). We considered at least one amplicon covering at least two gRNAs per gene as the minimum required for a knockout experiment and determined the fraction of genes per family that were ‘retained’ with these criteria for each design setting (here called ‘retention rate’).

**Figure 2:**
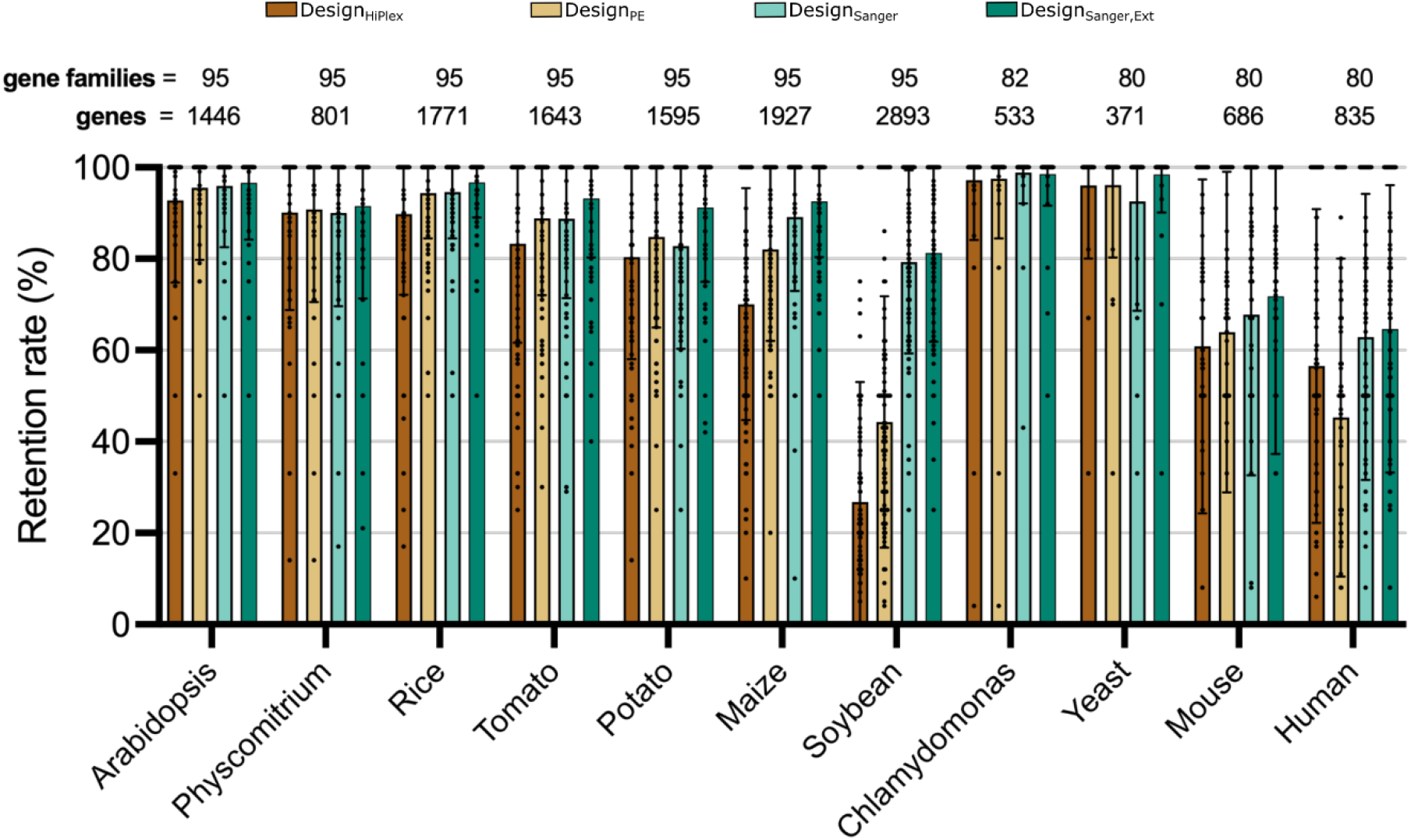
*SMAP Design* average retention rate across settings and species. Four Design settings were tested: Design_HiPlex_ (120 – 150 bp amplicons), Design_PE_ (220 – 250 bp), Design_Sanger_ (400 – 800 bp) Design_Sanger,Ext_ (400 – 800 bp with gene sequences extended by 500 bp at both ends). The same gene families were tested between settings and between the multicellular plant species and between the non-plant species, respectively. Retention rate is defined as the percentage of genes per gene family that contain at least one amplicon covering a minimum of two gRNAs. The bars show the average (with standard deviation) retention rate across all gene families per species.

Overall, the average retention rate per gene family for most tested species is ≥80% and there is a clear increase in retention rate with increasing amplicon size (**Figure 2**). The unicellular species Chlamydomonas and yeast had the highest average retention rates of ≥96%. Plant genomes with a lower fraction of recently-duplicated regions such as Arabidopsis, *P. patens*, and rice displayed average retention rates of 90% or higher for all three designs. The average retention rates for tomato, potato, and maize ranged from 80% to 90%, with the exception of maize for Design_HiPlex_ (65%). Soybean had the lowest average retention rate per gene family with 27%, 44%, 79% for Design_HiPlex_, Design_PE_ and Design_Sanger_, respectively, likely due to the highly duplicated nature of its genome (31, 32). These data indicate that genome constitution (e.g., recent genome duplication) affects the retention rate and that increasing the amplicon size can reduce design dropout.

During the analysis of Design_Sanger_, we also observed that small genes preferentially dropped out since the gene size was less than the required amplicon length of 400 – 800 bp. To overcome this limitation, *SMAP target-selection* was set to extract flanking regions 500 bp upstream and downstream of the target genes, resulting in Design_Sanger,Ext_. This increased the number of retained genes from 1399 to 1406 (out of 1446) genes for Arabidopsis (1% improvement), and from 1214 to 1338 (out of 1595) genes for potato (8% improvement; **Figure 2**). This further shows that primer and gRNA design optimization relies on both accurate selection of target reference sequences with *SMAP target-selection* as well as parameter settings of *SMAP design*.

The average retention rates were much lower for the mouse and human genomes (ranging from 45% to 68%). We reasoned that many of the 150 amplicons designed by default possibly fell into the relatively large introns and were thus filtered out. To overcome this, the option --*restrictedPrimerDesign* (*-rpd*) was added to restrict Primer3 to design amplicons near exons (**Supplementary Table S2**). Running *SMAP design* with *-rpd* increased the average retention rate for Design_PE_ by 10% for both species and the runtime was two to three times faster (**Supplementary Figure S3**). While ignoring intronic regions results in a clear improvement in retention rate, this result indicates there are additional sequence constraints limiting designs in these genomes and that parameter settings may need to be fine-tuned accordingly.

### Empirical testing of *SMAP design* in various species

To validate the specificity and amplification efficiency of Design_HiPlex_, we designed 40 amplicons on the *MAP3K* gene family in both Arabidopsis and soybean and performed HiPlex on 24 replicates of the reference genotypes for both species. For both Arabidopsis and soybean, all amplicons were sequenced in all replicates. Read depth was uniform across all amplicons with an average range <13-fold for 39 of the 40 amplicons for both species (**Figure 3**). Three amplicons for Arabidopsis and one amplicon for soybean fall below an arbitrary threshold of 1,000 reads across all samples and would likely be removed when establishing the genotyping assay to ensure reliable coverage of all amplicons in a screen. In Arabidopsis, a non-reference haplotype with a 1-bp deletion was found at the AT2G35050 locus with an average relative read depth of 3.8% across all samples. This deletion occurred in a homopolymer of ten adenosines and is therefore likely a sequencing error. Only the reference haplotypes were found for all other loci. The AT3G50730 amplicon displayed the lowest average read depth and failed for the five samples with the lowest overall total reads per library. In soybean, three loci displayed non-reference haplotypes. Two haplotypes contained two mismatches (SNPs) compared to the reference and the third haplotype had a 3-bp deletion compared to the reference in a “TCC” short sequence repeat. The average relative read depth of the non-reference haplotypes were consistently at or below 5% across all samples and can likely be attributed to low abundance PCR artifacts and/or sequencing errors. Such systematic errors can be removed with the haplotype frequency filters in *SMAP haplotype-window* or inclusion of the non-reference haplotypes in the FASTA reference sequence used for read mapping.

**Figure 3:**
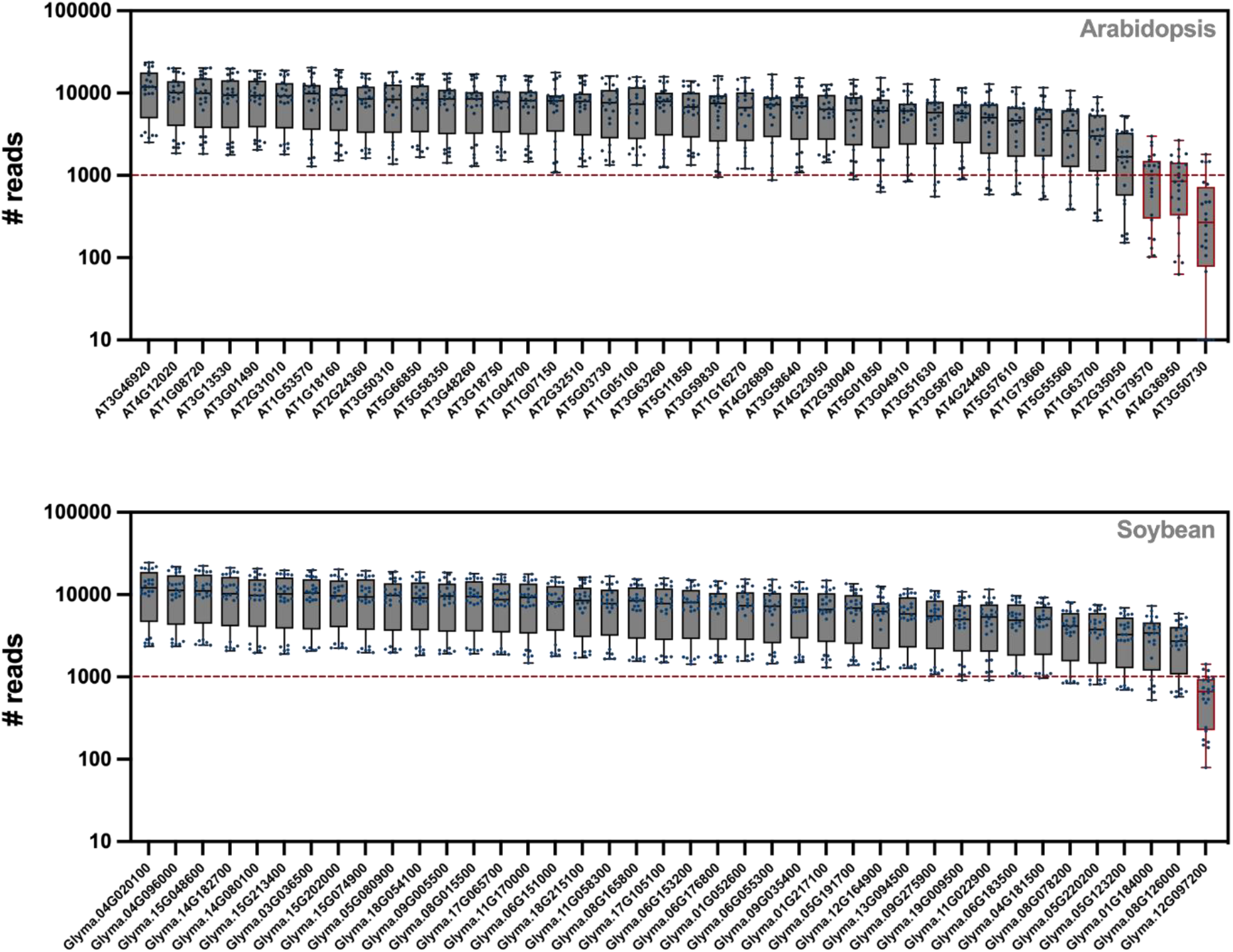
Read count per amplicon: For both the Arabidopsis (top) and soybean (bottom) genome, 40 amplicons were selected (one per gene) and sequenced using HiPlex sequencing on 24 replicate reference samples. The read depth per amplicon is given. The gene identifiers for which gene each amplicon was designed are given on the x-axis. The red dashed line indicates the desired minimum average read depth. Red boxes indicate amplicons with lower average read depth (these amplicons would be discarded and/or re-designed).

### Genome-wide *SMAP design*

As these initial experiments already generated designs for thousands of genes, we decided to pre-compute amplicons and gRNAs across the entire Arabidopsis genome as a resource for the research community. The Arabidopsis Col-0 reference genome contains 27,655 annotated genes assigned to 9,929 gene families containing between 1 and 208 genes (20). As we expected the potential for mispriming to be highest between genes from the same gene family, gene families were kept intact and divided into 28 groups of ±1000 genes to reduce the runtime via parallelization. *SMAP design* was run on each group to generate amplicons with Design_HiPlex_, Design_PE_, and Design_Sanger,Ext_. A maximum of three non-overlapping amplicons covering a maximum of two gRNAs per amplicon were designed per gene. The CPU runtime per group of ±1000 genes ranged from 16 to 78 hours (average 47 hours) on a server with Intel Xeon CPUs using one core per group. The retention rate (defined here as the fraction of genes within the gene family with at least one amplicon with at least one gRNA per gene) was ∼62% (17,186), ∼68% (18,886), and ∼85% (23,384) of all 27,655 Arabidopsis genes with Design_HiPlex_, Design_PE_, and Design_Sanger,Ext_, respectively.

In an effort to capture a greater fraction of the genes from the genome-wide designs, we checked if there were certain features that led to the dropout of amplicons and/or gRNAs. In particular, we wondered if genes from large gene families were preferentially filtered out in the gRNA/amplicon filtering steps, with the expectation that larger gene families would be overrepresented in the dropout gene set. We compared the relative distribution of the gene family size of the dropout genes to the genome-wide gene family size distribution and found an equal proportion of dropout genes across different gene family sizes (**Supplementary Figure S4;** two-sided Kolmogorov-Smirnov test *p*-value=0,87), suggesting that the retention rate is not biased towards a particular gene family size. We therefore questioned if the dropout was due to random matches of primers to non-gene-family members instead of sequence similarity to gene family members. To test this and increase the coverage of the genome-wide designs, a second run (here termed the dropout-only run) was performed where the dropout genes were run again but grouped only with other members of their gene family to still avoid potential mispriming between gene family members. Adding the designs from the dropout-only run increased the total coverage for Design_HiPlex_, Design_PE_, and Design_Sanger,Ext_ to ∼92%, ∼94%, and ∼96%, respectively. Thus, the dropout genes were likely lost due to Primer3-predicted non-specific primer binding onto reference sequences outside of their gene families.

A similar two-step approach was followed for the *P. patens* genome where the 32,926 genes were divided into 33 groups of ±1,000 genes, keeping gene family members together (20). The genome consists of 15,604 gene families with 1 to 273 members. The Design_Sanger,Ext_ run yielded amplicons and gRNAs for ∼77% of the genome and was increased to ∼86% by including the designs from the additional dropout-only run. The CPU runtime for a group of ± 1,000 genes ranged from 26 to 146 hours (average 113 hours).

To validate the Arabidopsis genome-wide Design_Sanger_,_Ext_ and check for off-target amplification, 48 primer pairs were selected from each of the first and dropout-only runs of the genome-wide design. PCR followed by gel electrophoresis showed that 94 out of 96 amplicons had a single visible band (**Supplementary Figure S5**) and the two other amplicons (one from the first run and one from the dropout-only run) showed a single, yet less intense band. High-quality Sanger sequencing reads were obtained for 85 out of the 96 PCR products and confirmed all amplicons specifically amplified one single locus. Overall, since no discrepancy was found between the first and dropout-only runs, either through gel electrophoresis or through Sanger sequencing, we conclude that the primers are efficient and specific for Sanger sequencing and our observations suggest that the Primer3 default settings for eliminating non-specific primers were too conservative for Sanger sequencing. We therefore added an option for users to adjust the specificity settings of Primer3 when running *SMAP design*.

The eleven low-quality reads contained stretches with ≥ 10 thymidines or adenosines (homopolymers) resulting in overlapping sequencing peaks which are problematic for Sanger and Illumina-based genotyping. Interestingly, homopolymers with < 10 repeated nucleotides were observed in the sequenced amplicons but did not lead to overlapping sequencing peaks. At least under our Sanger sequencing conditions, there appears to be a threshold of 10 repeated nucleotides. We therefore calculated how many potentially problematic homopolymers were present in the amplicons of genome-wide design of Design_Sanger,Ext_ for Arabidopsis. Out of the 45,189 amplicons, 2,889 (6.4%) had at least one homopolymer (≥ 10 nucleotides) with the majority of the homopolymers consisting of poly-A or poly-T (>99%) (**Supplementary Figure S6**). A filter was therefore implemented in *SMAP design* to remove amplicons containing homopolymers of a user-defined size (**Supplementary Table S2**). Running *SMAP design* on Design_Sanger,Ext_ with the homopolymer filter (-*hp*) set to 10 nucleotides for the genome of Arabidopsis and *P. patens* yielded a retention rate of ∼95% and ∼85% respectively (including the dropout-only runs). These final genome-wide designs are available in Supplementary data.

### Using *SMAP design* to screen for natural variation

#### Amplicon design and detection rates

To evaluate the use of *SMAP design* to screen for natural variation in a non-model organism (chicory), HiPlex amplicons were designed to sequence nine candidate genes putatively involved in haploid induction (33–35). We aimed to create a catalogue of naturally-occurring sequence variants and ideally find haplotypes affecting the protein sequences as these could be used to generate haploid-inducer lines. We created two complementary HiPlex primer sets with *SMAP design* using Design_NatVar_ settings that contain partially overlapping (tiled) amplicons for each of the nine genes for a total of 94 amplicons (**Figure 4**). We screened 35 chicory (*C. intybus* var. *sativum*) and 25 witloof accessions (*C. intybus* var. *foliosum*) by applying a 1-D pooling strategy (**Supplementary Figure S7A**) in which an equal amount of leaf material from ∼10 individuals was pooled for a single DNA extraction and three independent pools per accession were created (n∼30 plants). In total, using the two HiPlex assays, 1,554 chicory plants were screened in 163 pools, and pools with interesting sequence variants were identified. Individual plants from selected pools were then sequenced to identify carriers of knockout alleles.

**Figure 4:**
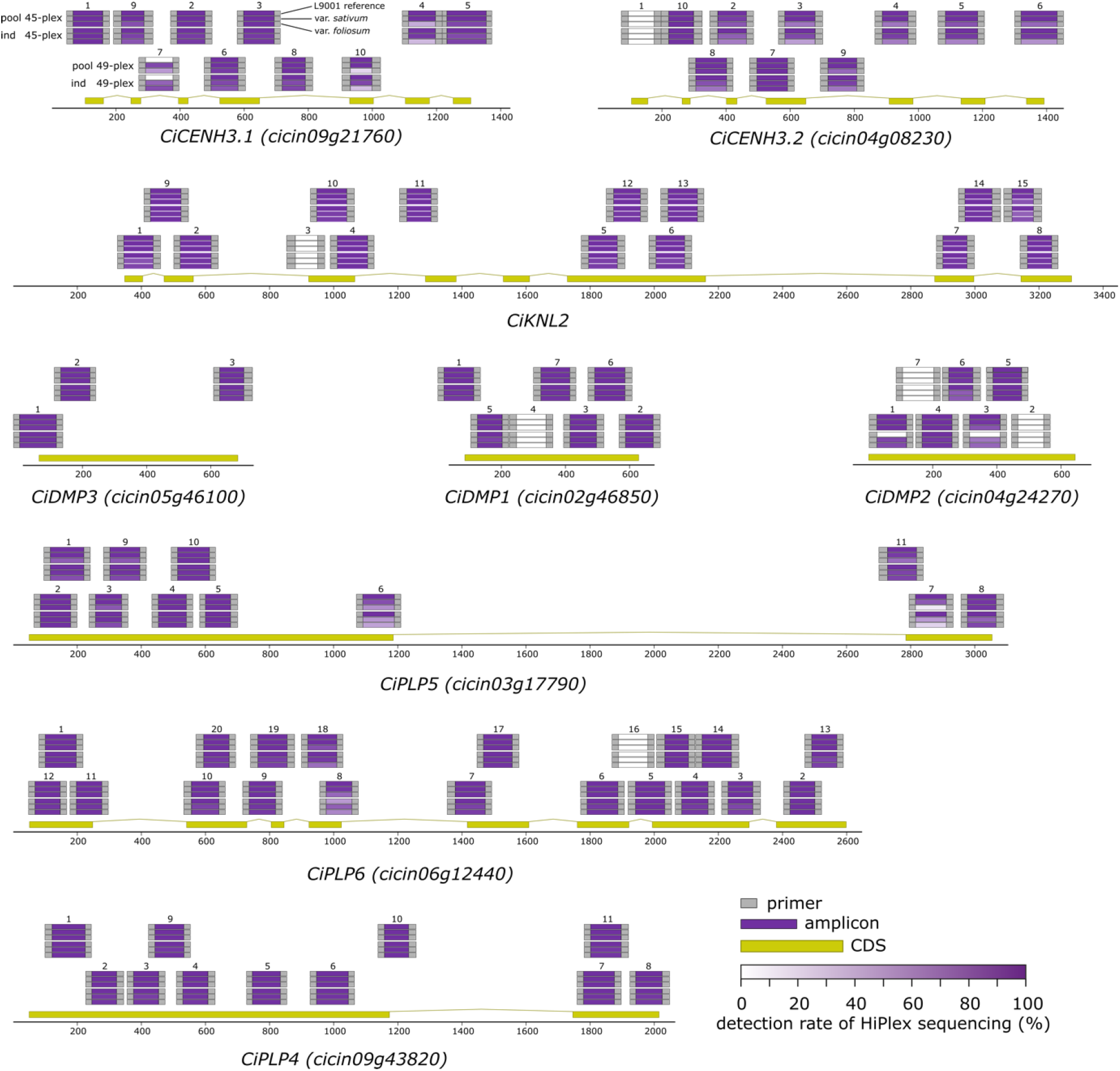
Detection of HiPlex amplification of two (tiled) amplicon designs for nine candidate genes putatively involved in haploid induction. Data from the 45-plex and the 49-plex assay in pooled (pool) and individual (ind) sequencing runs are included. For each amplicon and each run, the detection rate is shown, split by replicates of the L9001 reference genotype (top bar), *C. intybus* var. *sativum* samples (middle bar), and *C. intybus* var. *foliosum* samples (lower bar).

We evaluated primer design performance by assessing the absolute read counts per amplicon. We set a “detection” threshold at a minimum of 30 reads per amplicon per sample and defined the “detection rate” per amplicon as the percentage of samples with read depth greater than 30 for that amplicon. Six amplicons were removed from further analysis because they either yielded no sequencing reads (CiDMP2_02, CiPLP6_16, and KNL2_03) or had a read depth below 100 reads across all samples (CiCENH3.2_01, CiDMP1_04, and CiDMP2_07). In the pooled sequencing run, 87 of the 88 remaining amplicons were detected in at least 90% of the reference samples (**Table 1**). The number of amplicons detected in at least 90% of the samples dropped in the individual sequencing run compared to the pooled sequencing run for both chicory and witloof, with chicory having a higher detection rate compared to witloof. For chicory, the detection rate dropped from 97% to 82%, and for witloof from 77% to 73% for pools and individuals, respectively (**Table 1**), confirming the expectation that amplification becomes less efficient with increasing genetic distance from the chicory reference genome sequence. Overall, we were able to cover 36% to 92% of the CDS per gene, after considering the design and amplification dropouts (**Figure 4, Table 1**).

#### Identification of conserved and variable gene regions in a breeding genepool

We used *SMAP haplotype-window* to list the number of different haplotypes per amplicon per sample and estimated the relative haplotype frequencies in the pooled dataset (**Supplementary Table S5**). We compared the overall number of haplotypes and haplotypes leading to protein changes between chicory and witloof accessions in the pooled dataset. In the chicory accessions, 257 different haplotypes were found across the 88 amplified loci, while for the witloof accessions 519 different haplotypes were found across all 88 loci, of which 242 haplotypes are found in both chicory and witloof, illustrating a higher level of sequence variation within these genes in the witloof accessions. In the chicory and witloof accessions, 93 (36%) and 247 (48%) of the haplotypes led to a different protein sequence, respectively. Of the 267 unique haplotypes found leading to protein changes across both chicory and witloof accessions, 253 (95%) were SNPs or in-frame mutations, while only 14 haplotypes (5%) were frameshift mutations. A total of 21 amplicons located in exonic regions did not have any haplotypes leading to protein changes and could thus be considered as conserved genic regions. The number of haplotypes per amplicon varied between genes, and across the length of the gene sequences (**Figure 5**). For instance, in *CiCENH3*.*1* and *CiCENH3*.*2*, more haplotypes and haplotypes leading to protein changes were found in the N-terminal region of the genes. The average number of all haplotypes per amplicon per gene ranged from 4.2 (*CiPLP4*) to 7.6 *(CiCENH3*.*1*), and the average number of haplotypes per amplicon per gene with protein sequence changes ranged from 1.2 (*CiCENH3*.*2*) to 4.4 *(CiKNL2*).

**Figure 5:**
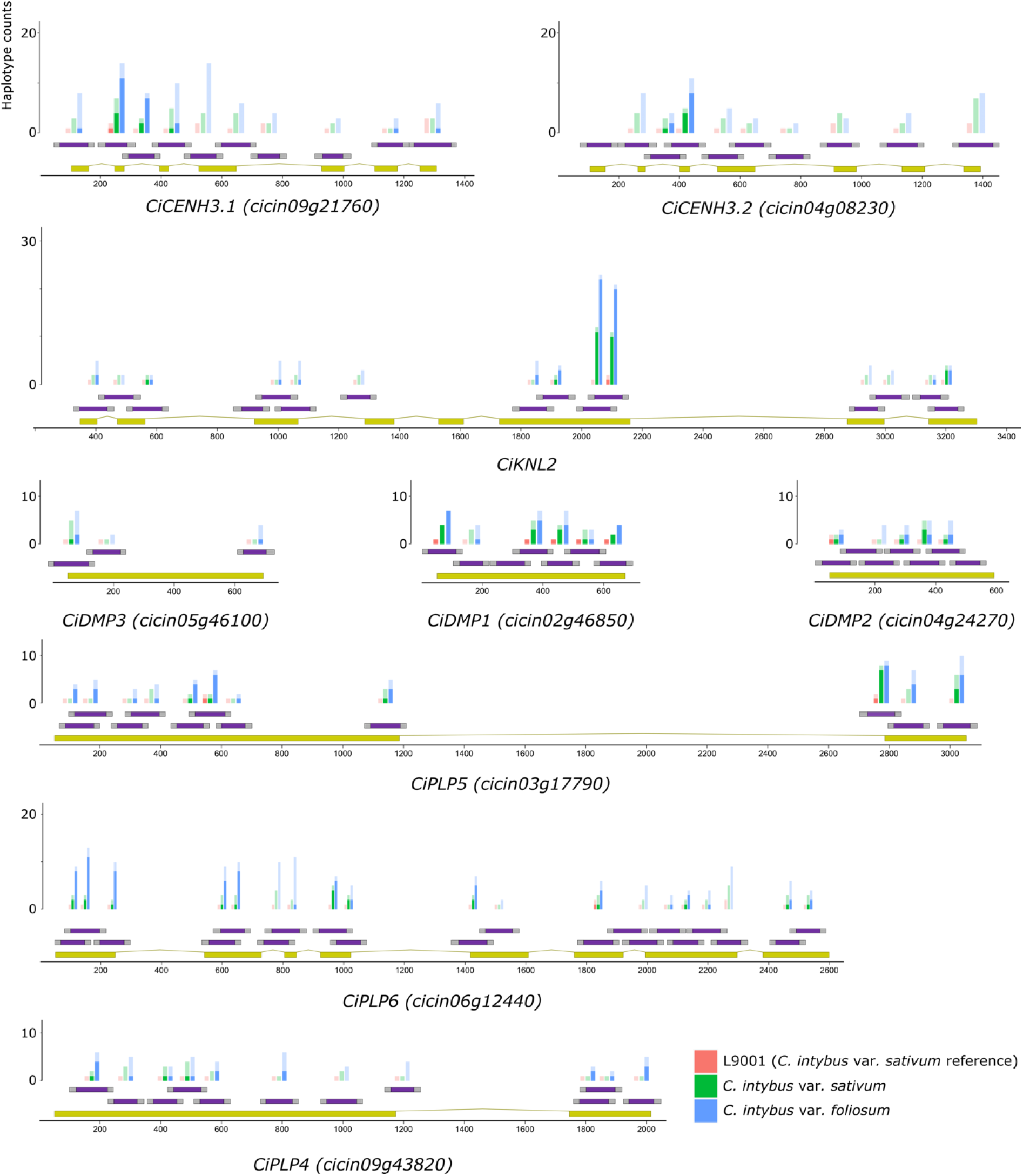
Observed haplotypes in *Cichorium intybus* pooled samples. The total number of haplotypes (transparent) and the number of haplotypes with protein sequence changes (opaque) per amplicon and per accession are indicated as bars. For the L9001 reference genotype, 6 replicates were included. For 35 accessions of *C. intybus* var. *sativum*, a total of 1039 plants were screened in 104 pools. For 25 accessions of *C. intybus* var. *foliosum*, a total of 465 plants were screened in 54 pools.

We focused on haplotypes with changes in the predicted protein sequences to identify individuals with potential knockout alleles, defined here as a protein sequence similarity of less than 50% compared to the reference protein sequence. Using *SMAP effect-prediction*, we predicted the effect of haplotypes on the protein function as “mild effect” if more than 50% of the resulting protein sequence was identical to the reference protein and as “strong effect” if at most 50% of the protein sequence was identical to the encoded reference protein. 255 haplotypes were classified as a mild effect on protein function (48% of all haplotypes) and 12 haplotypes were classified as strong effect (2% of all haplotypes; **Table 2, Supplementary Table S5**). 320 plants (from 36 initial pools, 15 from chicory, 21 from witloof) with strong-effect haplotypes were sequenced individually. Due to the loss of plants between sampling of the pools and individuals, data were obtained for 294 individuals (91%). We recovered a total of 13 strong-effect haplotypes in *CiPLP6, CiPLP5, CiDMP1*, and *CiDMP2. CiPLP6* was the most variable, with 10 strong-effect haplotypes, which were found 84 times in a heterozygous state and eight times in a homozygous state across 46 individuals and were often combined in a single individual (**Table 2**). Strong protein effect haplotypes in *CiPLP5* (2 individuals), *CiDMP1* (2 individuals), and *CiDMP2* (1 individual) were all present in a heterozygous state (**Table 2**).

**Table 2:** Effects of mutations on predicted protein sequence from haplotypes detected in pools and individual (ind) sequencing. Haplotypes with an effect on the protein sequence were defined as ‘mild’ effect if more than 50% of the resulting protein sequence was identical to the reference protein, or as ‘strong’ effect if at most 50% of the protein sequence was identical. Abbreviations: HE = heterozygous, HO = homozygous. NA* The gene identifier for this gene is missing in the latest annotation of the genome.

| Gene | GeneID | CDS coverage (%) | Total number of haplotypes (pools / ind) | # haplotypes mild effect (pools / ind) | # haplotypes strong effect (pools / ind) | # carriers identified HE / HO (pools) |
| --- | --- | --- | --- | --- | --- | --- |
| *CiCENH3.1* | *cicin09g21760* | 92.27 | 76 / 71 | 23 / 21 | 0 / 0 | 0 / 0 (0) |
| *CiCENH3.2* | *cicin04g08230* | 79.73 | 51 / 43 | 11 / 5 | 0 / 0 | 0 / 0 (0) |
| *KNL2* | NA* | 65.40 | 96 / 65 | 60 / 39 | 1 / 0 | 0 / 0 (0) |
| *CiDMP1* | *cicin02g46850* | 79.23 | 32 / 27 | 27 / 18 | 3 / 1 | 2 / 0 (2) |
| *CiDMP2* | *cicin04g24270* | 70.17 | 22 / 20 | 11 / 6 | 0 / 1 | 1 / 0 (1) |
| *CiDMP3* | *cicin05g46100* | 35.66 | 13 / 11 | 3 / 3 | 0 / 0 | 0 / 0 (0) |
| *CiPLP5* | *cicin03g17790* | 57.29 | 62 / 48 | 40 / 28 | 1 / 1 | 2 / 0 (1) |
| *CiPLP6* | *cicin06g12440* | 80.53 | 136 / 134 | 63 / 59 | 7 / 10 | 84 / 8 (16) |
| *CiPLP4* | *cicin09g43820* | 58.64 | 46 / 46 | 17 / 17 | 0 / 0 | 0 / 0 (0) |
| Total |  |  | 534 / 465 | 255 / 196 | 12 / 13 | 89 / 8 (20) |

#### Accuracy and sensitivity of haplotype detection in pool-Seq

With the pooled and individual sequencing data, we assessed the sensitivity of pooled sequencing combined with HiPlex to identify haplotypes across a range of candidate genes in parallel. We analyzed the data from 13 pools of 10 individuals with complete sequencing data in all 130 individuals (ground truth) and calculated the ‘expected’ pooled haplotype frequency based on the discrete genotype calls of the 10 constituent diploid individuals. For instance, a single heterozygous individual in a pool of 10 diploid plants (i.e., 20 alleles) corresponds to a 5% relative haplotype frequency in HiPlex pooled sequencing data. A strong correlation (R² value: 0.8487) was found between the observed haplotype frequencies in pools and the haplotype frequencies in individuals (**Figure 6A**). Additionally, 1,826 of all 1,933 (94.5%) haplotypes detected in individuals were also detected in their respective pools (true positives in pools), and 107 haplotypes (5.5%) were only detected in the respective individuals (false negatives in pools). About half (59/107) of the false-negative haplotypes displayed an ‘expected’ haplotype frequency of 5% or 10% in the individual sequencing data of the respective pool, indicating that pooled sampling effectively detects almost all haplotype diversity, with a weak bias against very low frequency haplotypes (**Figure 6B**). Conversely, 106 of 1,932 (5.5%) haplotypes detected across all pools were not detected in their respective constituent individuals (false positive in pools). The observed haplotype frequencies of 89 of 106 (84.0%) of the false positives were in the range of 1-5%, characteristic of low frequency read errors (**Figure 6B**). Taken together, these data show that HiPlex pooled sequencing (n=10) accurately quantifies the relative frequency of haplotypes within a pool and is sensitive enough to detect nearly all low-frequency haplotypes, including rare defective alleles.

**Figure 6:**
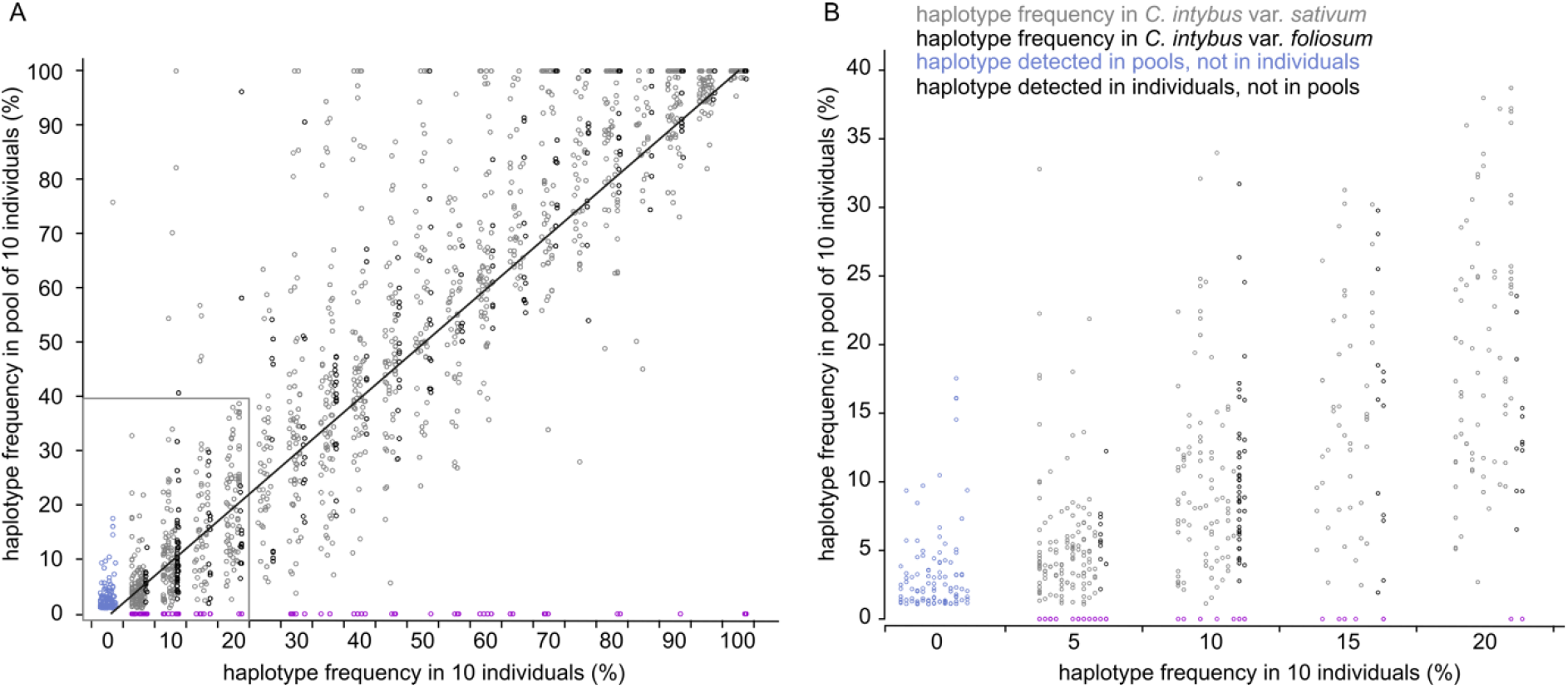
Comparison of haplotype frequency in pools to haplotype frequency in their 10 constituent individuals. **A)** The diagonal line shows the expected ratio where the observed PoolSeq haplotype frequency is equal to the expected haplotype frequency based on individual sequencing. **B)** Enlargement of the small panel inset in A.

## DISCUSSION

There is an increasing need for scalable PCR amplicon design for genotyping as the scale of eco-tilling and multiplex CRISPR experiments continues to increase. This is particularly the case when conducting CRISPR screens with at least ∼50 genes as manual designs can take several weeks, if not months, to perform. The currently-available CRISPR design tools can generate lists of genome-wide gRNAs, but they lack the ability to combine this with genotyping primer design in a flexible and customizable manner. Tools such as CHOPCHOP (14), CRISPOR (13), or CROPSR (36) allow gRNA and associated primer design, but the user is very limited in the ability to customize it (e.g., number of gRNAs per amplicon, number of amplicons per gene, relative position of gRNAs along the gene, etc.). In addition, these tools provide the user with tens to hundreds of designs per gene, leaving them to sort through the designs that are compatible with each other in a multiplex format. *SMAP design* overcomes these limitations by designing highly specific amplicons at gRNA target sites for any number of user-selected genes and presents the user with compatible, ready-to-order designs. *SMAP design* greatly simplifies larger CRISPR experiments (e.g., multiplex and combinatorial CRISPR screens (37)) by reducing the design step from weeks or months to just hours of CPU time. The pre-designed genome-wide amplicons presented here effectively eliminate the design step altogether as users just need to select their gene identifier in the list and order the associated primers and/or gRNAs. Based on our empirical tests using HiPlex and Sanger sequencing, these pre-computed amplicons are reliable and amplify the target loci with high specificity. The resulting Sanger or NGS data can then be seamlessly analyzed with ICE (22), TIDE (23), or other tools from the SMAP package (19) using the output files from *SMAP target-selection* and *SMAP design* as input (**Figure 1**).

Similarly, *SMAP design* can be used to screen for naturally-occurring sequence variants with high specificity and sensitivity. For eco-tilling applications, amplicon design relies on gene-specific primers, limited off-target amplification, and covering as much of the gene sequence as possible. Current amplicon design methods consist of primer design by Primer3, followed by a BLAST and/or a preliminary PCR to check for mispriming (38). This can become laborious and time-consuming for large numbers of target genes. *SMAP design* automates this amplicon design and avoids mispriming. Amplicon design for the nine candidate genes in chicory proved to be reliable and gene specific and amplification was robust in different genotypes and accessions, thus capturing a broad range of sequence variation across the breeding genepool.

While the number of genes that *SMAP design* can handle is theoretically unlimited, it is not always possible to design gene-specific amplicons or gRNAs for all genes. As shown here, genomes with relatively recent whole genome duplications or polyploid genomes suffer from lower retention rates, most likely due to the primer specificity checks implemented in Primer3. The retention rate depends on the genome and gene family but is generally higher than 80% for most of the tested species. Generating designs in species with highly duplicated genomes, such as soybean, can be more challenging. We show that increasing amplicon size, restricting primer design to exons and splitting up gene families or groups can increase the retention rate. Furthermore, relaxing the default primer specificity settings in Primer3 would likely increase the retention rate as well. The settings that can be changed are of course dependent on the type of screen that will be performed; if simplex PCR will be used (e.g., for Sanger sequencing), cross-amplification is of no concern so relaxing the primer-specificity filters can be tolerated, but cross-amplification would be problematic for highly multiplex PCR where mixtures of primer pairs are used. As there are a wide range of settings, options, and variables (genotyping assay, genome, and target genes), we recommend the practical approach is to first empirically validate all genotyping assays by sequencing reference genotypes and eliminate any primers or amplicons that do not amplify efficiently or are non-specific. This will ensure a smooth genotyping workflow once the mutant materials are generated and need to be characterized. This also ensures that there are no sequence variants in the gRNA targets between the reference sequence used for the design and experimental genotypes, and if there are, corrections can be made before cloning is initiated. Since the composition of the reference sequence influences the specificity of primer and gRNA designs (and thus the overall design retention rate), *SMAP target-selection* is an important utility tool to streamline the construction of alternative reference gene sets and customize input parameter settings for optimal retention rate and reference sequence coverage, while maintaining target specificity.

Some limitations to designing gRNAs and amplicons in a high-throughput fashion remain. For instance, it is not yet possible to design gRNAs and corresponding amplicons that target multiple genes at the same time. Programs such as CRISPys (39) or MultiTargeter (40) can design gRNAs that allow the targeting of multiple loci by exploiting the capability of gRNAs with mismatches to the target sequences to still be functional. A gRNA list from such a program could be fed to *SMAP design*, however, amplicons will only be designed for the targets with an identical gRNA sequence and primers binding multiple regions will be filtered out. Furthermore, the on-target efficiency scores such as Doench (29) and Out-of-Frame (13) have not translated well to plants (41). Therefore, it is not guaranteed that a gRNA in the output of *SMAP design* will result in a knockout even though it might have a high efficiency prediction. Efficiency scores trained on plant data are thus highly desirable.

We also illustrated the capacity for HiPlex amplicon sequencing to perform eco-tilling by detecting low frequency alleles (5%) in a pool of 10 individuals. Indeed, to sequence very large numbers of individuals, 1-D pooling of 10 plants per pool substantially reduces the time, effort, and cost for DNA extraction, library preparation and sequencing, while retaining detection accuracy and sensitivity for low frequency haplotypes (**Supplementary Figure S7A**). Screening efficiency can be further increased with 2-D or 3-D pool sequencing approaches, routinely used in tilling by sequencing applications (38). For example, a 2-D pooling scheme based on a pool size of 10 plants divides 100 plants into 2 × 10 = 20 pools, each containing 10 plants in which each plant becomes part of 2 pools (with X_1-10_ and Y_1-10_ coordinates; **Supplementary Figure S7B**). This leads to a 5-fold reduction in the number of PCRs (20 PCRs on pools instead of 100 PCRs on individuals). Given that we observe a minimum detection threshold of ∼5% allele frequency per pool, this allows us to detect a single heterozygous mutation in a pool of 10 plants. If a particular mutation occurs only once in the set of 100 (diploid) plants, the plant carrying the mutation can be identified in the corresponding X- and Y-pools. If the same mutation occurs more than once in the set of 100 plants (>1% population frequency), a second round of screening at the individual plant level is required at each of the intersecting X- and Y-pool coordinates. As knockout alleles are quite rare, natural variants, such a 2-D or even 3-D pooling approach can be used to quickly identify such alleles and their carriers in a cost-effective manner. Combined with HiPlex amplicon sequencing, which can screen multiple loci and genes at once, this allows for rapid screening of many genes in large populations. While we demonstrated the versatility of pooled sequencing to screen for natural variation, it is clearly a useful strategy for screening large collections of CRISPR mutants, as each unique type of mutation (defined by the haplotype sequence) is identified independently.

Ultimately, we envision a reverse-genetics approach where a researcher would use *SMAP design* to first screen their gene pool material for natural knockout alleles, as demonstrated for *CiDMP1, CiDMP2, CiPLP5*, and *CiPLP6*. The identified carriers of the alleles could then be utilized for functional analysis and/or breeding. Alternatively, if no genetic variation is found, the researcher then resorts to induced genetic mutations where CRISPR is a highly tractable option for transformable species. For example, we did not observe any strong-effect haplotypes in *CiCENH3*.*1, CiCENH3*.*2, CiKNL2, CiDMP3*, or *CiPLP4*. Therefore, the straightforward way to continue investigating these genes for haploid induction is to utilize CRISPR mutagenesis of chicory (42). The target sequences have been confirmed in our genepool material via HiPlex, and the overlapping gRNAs can directly be cloned into Cas9/gRNA expression vectors. Overall, *SMAP design* will be a useful tool to perform high-throughput genetic screens using both natural and induced variation.

## AVAILABILITY

All tools within the SMAP package (*SMAP target-selection, SMAP design, SMAP haplotype-window, SMAP effect-prediction*) and pre-computed designs are available in the GitLab repository https://gitlab.com/ilvo/smap-design and https://gitlab.com/truttink/smap. Manuals can be found at https://ngs-smap.readthedocs.io/.

FlashFry and CRISPOR for gRNA design can be found at https://github.com/mckennalab/FlashFry and https://github.com/maximilianh/crisporWebsite respectively.

PLAZA (https://bioinformatics.psb.ugent.be/plaza/) was used to retrieve genomes and annotations files.

BWA-MEM (https://github.com/lh3/bwa) was used to map reads to the reference sequences.

*SMAP target-selection* and *SMAP design* are also being made available via Galaxy at https://usegalaxy.be/.

Amplicon NGS files were deposited at SRA (https://www.ncbi.nlm.nih.gov/sra) under the accession numbers PRJNA848638 (Arabidopsis and soybean) and PRJNA855321 (chicory).

## SUPPLEMENTARY DATA

Supplementary Data are available at NAR online.

## ACKNOWLEDGEMENT

We thank the ELIXIR Belgium team (supported by Research Foundation-Flanders, project I002819N) for the assistance in making the tool available in Galaxy. We thank Dries Schaumont for the guidance with the NGS analysis.

