## Supplementary Tables and Figures for "*SMAP design*: A multiplex PCR amplicon and gRNA design tool to screen for natural and CRISPR-induced genetic variation"

**Table S1: Genomes used in this study for each species.** All genome sequences and annotations are available at https://bioinformatics.psb.ugent.be/plaza/. Dicots PLAZA 4.5, Monocots PLAZA 4.5, and Pico-PLAZA 3.0 were used to retrieve all genomes. An unpublished, in-house assembled, and annotated reference genome sequence of *Cichorium intybus* var. *sativum* was used for genome-wide identification of gene family members from the selected gene families.

| **Species** | **Common name** | **Genome version** |
|---|---|---|
| *Arabidopsis thaliana* | Arabidopsis | Araport11 |
| *Glycine max* | Soybean | JGI Wm82.a2.v1 |
| *Oryza sativa ssp. japonica* | Rice | IRGSP-1.0-2021-05-10 |
| *Physcomitrium patens* | Physcomitrium | JGI v3.3 |
| *Solanum lycopersicum* | Tomato | Sol Genomics itag2.4 |
| *Solanum tuberosum* | Potato | JGI v4.03 |
| *Zea mays* | Maize | NAM v5.0 |
| *Chlamydomonas rheinhardtii* | Chlamydomonas | JGI v5.5 |
| *Saccharomyces cerevisiae strain S288C* | Yeast | R64-1-1 |
| *Mus musculus* | Mouse | GRCm38.p4 |
| *Homo sapiens* | Human | GRCh38.p2 |
| *Cichorium intybus* var. *sativum* | Chicory | unpublished |

**Table S2: Optional parameters for *SMAP design*.** Full and abbreviated names of the user-defined parameters in *SMAP design* and their actions. The default settings are indicated in square brackets.

| Argument | Abbr. | Description |
|---|---|---|
| **Input options** | | |
| --gRNAfile | -g | gRNA file (tsv). |
| --gRNAsource | -gs | The source of the gRNA file: "CRISPOR", "FlashFry" or "other" [**FlashFry**]. |
| --preSelectedPrimers | -psp | Set of primers/amplicons (GFF) for which gRNAs should be found. |
| --selectGenes | -sg | List of genes to which amplicons and gRNAs must be designed. The other genes in the FASTA file will be used to check for specificity only. |
| **Amplicon options** | | |
| --generateAmplicons | -ga | Number of amplicons to generate per gene by Primer3 [**150**]. |
| --restrictedPrimerDesign | -rpd | Exclude Primer3 from generating primers in large introns. |
| --minimumAmpliconLength | -minl | The minimum length of the amplicons in base pairs [**120**]. |
| --maximumAmpliconLength | -maxl | The maximum length of the amplicons in base pairs [**150**]. |
| --primerMaxLibraryMispriming | -pmlm | The maximum allowed weighted similarity of a primer with any sequence in the target gene set (Primer3 specificity setting) [**12**]. |
| --primerPairMaxLibraryMispriming | -ppmlm | The maximum allowed sum of similarities of a primer pair (one similarity for each primer) with any single sequence in the target gene set (Primer3 specificity setting) [**24**]. |
| --primerMaxTemplateMispriming | -pmtm | The maximum allowed similarity of a primer to ectopic sites in the template (Primer3 specificity setting) [**12**]. |
| --primerPairMaxTemplateMispriming | -ppmtm | The maximum allowed summed similarity of a primer pair to ectopic sites in the template (Primer3 specificity setting) [**24**]. |
| --homopolymer | -hp | The minimum number of repeated identical nucleotides in an amplicon for the amplicon to be discarded [**10**]. |
| --misPrimingAllowed | -mpa | Do not check for mis-priming when designing primers. |
| --numberAmplicons | -na | The maximum number of non-overlapping amplicons per gene [**2**]. |
| --ampliconLabel | -al | Number the amplicons from left to right. |
| **gRNA filtering options** | | |
| --promoter | -prom | The last 6 bases of the promoter for the gRNA to check for BsaI or BbsI sites in the promoter-gRNA-scaffold sequence [**TGATTG**]. |
| --scaffold | -scaf | The first 6 bases of the scaffold to check for BsaI or BbsI sites in the promoter-gRNA-scaffold sequence [**GTTTTA**]. |
| --gRNAOverlap | -go | The minimum number of bases between the start of two gRNAs in an amplicon [**5**]. |
| --threshold | -t | Minimum gRNA specificity score (MIT) [**80**]. |
| --restrictionSite | -rs | Do not filter out gRNAs that contain a BsaI or BbsI restriction site. |
| --targetRegion5 | -tr5 | The fraction of the coding sequencing at the 5' end that is excluded from being targeted (expressed in percentage of total CDS length) [**0.2**]. |
| --targetRegion3 | -tr3 | The fraction of the coding sequencing at the 3' end that is excluded from being targeted (expressed as percentage of total CDS length) [**0.2**]. |
| --distance | -d | The minimum number of bases between primer and gRNA [**15**]. |
| --numbergRNAs | -ng | The maximum number of gRNAs to retain per amplicon [**2**]. |
| --gRNAlabel | -gl | Number the gRNAs from left to right. |
| --targetSpecificRegion | -tsr | Only target a specific region in the gene indicated by the feature name in the GFF file. |
| --polyT | -pT | Minimum number of repeated Ts (in a poly-T) in the gRNA to avoid [**4**]. |
| **Output options** | | |
| --output | -o | Name of the output files [**SMAPdesign**]. |
| --allAmplicons | -aa | Return all amplicons with their respective gRNAs after filtering per gene. |
| --borderLength | -b | The length of the borders (for *SMAP haplotype-window*) [**10**]. |

| | | |
|---|---|---|
| --borderOnly | -bo | Write additional GFF file with only borders (for *SMAP haplotype-window*), or a BED file with SMAP sites (for *SMAP haplotype-sites*. |
| --debug | -db | Gives a GFF file with all amplicons designed by Primer3 and all gRNAs before filtering. |
| --summary | -smy | Write summary table and plot graphs of the output. |
| --verbose | -v | Report progress of primer design. |

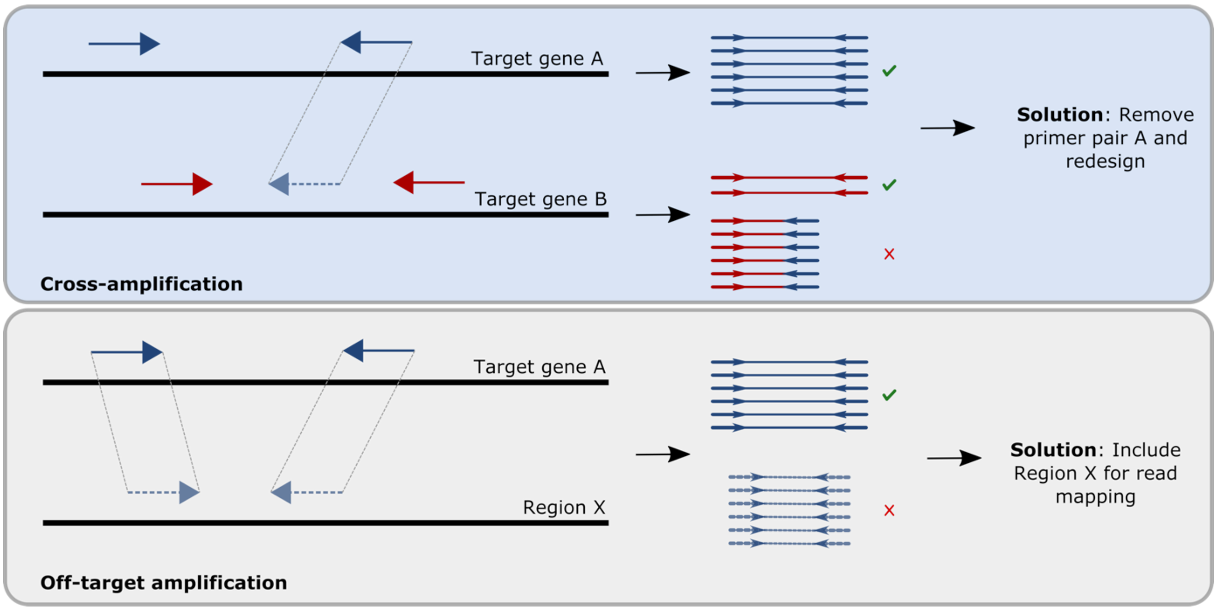


**Figure S1. Examples of mispriming errors during PCR-based genotyping.** In multiplex PCR, cross-amplification is the case where one or more primers from another amplicon (A) bind in between a primer pair of a given amplicon (B). This results in a truncated PCR product and incomplete coverage of the amplicon (B). The solution is to remove the problematic primer and redesign the assay. Off-target amplification is the case where primers of gene A bind elsewhere in the genome (Region X) and results in two unique amplicons. In this case, the extra amplicon can be excluded from mapping at location target gene A during read mapping analysis by including region X into the reference sequence.

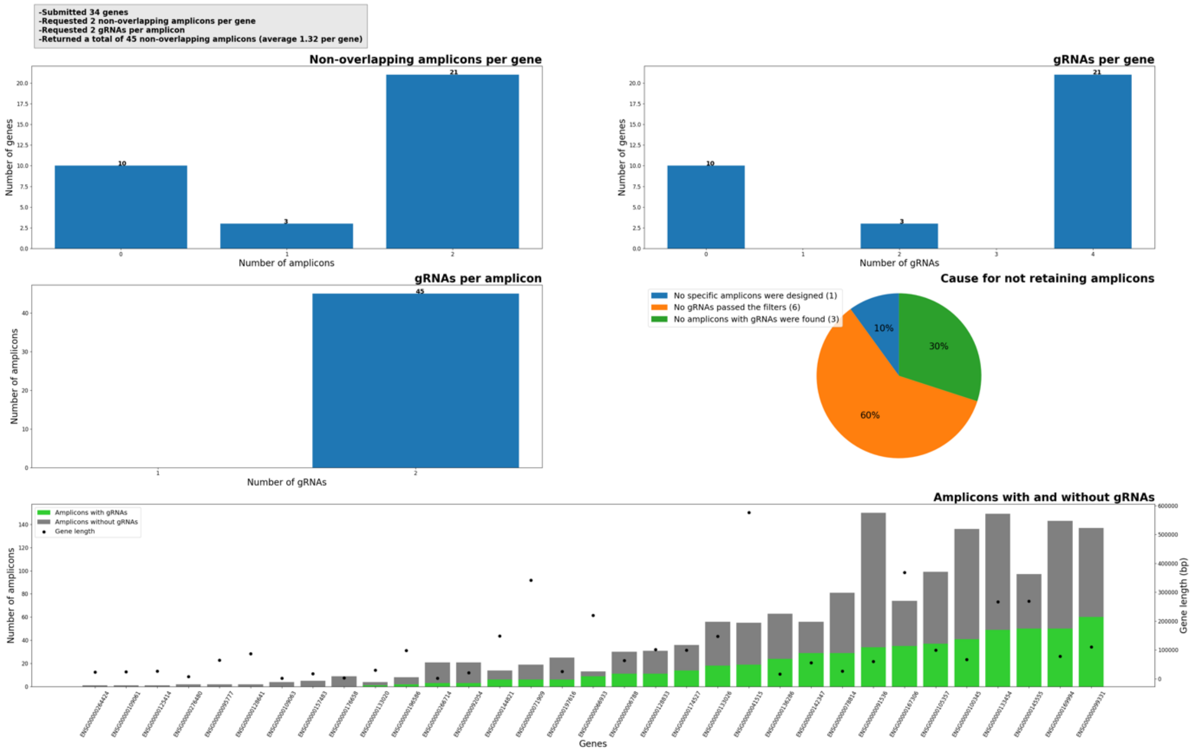


**Figure S2: Summary graph optionally created by *SMAP design*.** The upper-left box shows information on the input from the user and summarizes the output. The upper-left plot shows the number of genes for which a certain number of amplicons have been designed. The upper-right plot shows the number of genes that retained a certain number of gRNAs. The middle-left plot shows the number of amplicons which cover a certain number of gRNAs. The middle-right plot shows the percentage of genes that were not retained and explains why no amplicons could be designed and/or retained. The bottom plot shows for each gene, the total number of amplicons that were designed by Primer3 and how many of these amplicons covered any gRNAs or covered no gRNAs. The dots show the total gene sequence length (in bp).

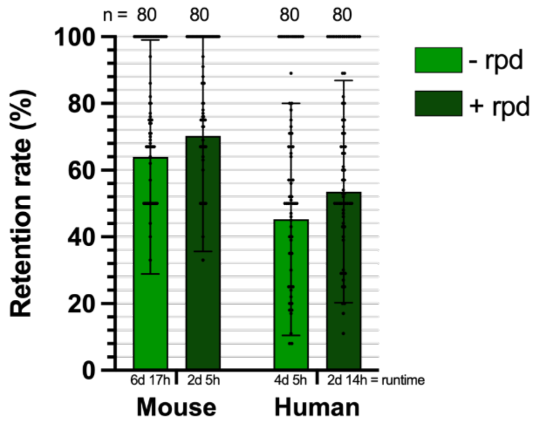


**Figure S3: Retention rate of Design$_{PE}$ with and without option rpd for mouse and human**. *SMAP design* was run with 80 gene families of the mouse and human genome with the *restrictedPrimerDesign (rpd)* function either on or off. The function limits Primer3 to only generate amplicons to exonic regions, avoiding large introns typical for the mouse and human genome. CPU runtime is indicated in days (d) and hours (h). – rpd, *SMAP design* was run with the rpd option off; + rpd, *SMAP design* was run with the rpd option on.

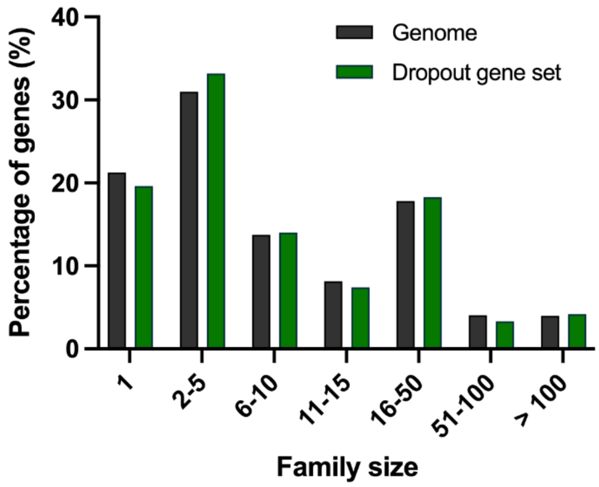


**Figure S4: Distribution of the gene family size of the dropout genes from the genome-wide Design$_{Sanger,Ext}$ and the whole genome.** *SMAP design* was run on all 27,655 genes grouped in 9,928 gene families in the Arabidopsis genome. No amplicons were retained for 4,269 genes, belonging to 2,218 gene families of one or more genes. The bars represent the relative number of genes belonging to a gene family of a particular size range, either in the whole Arabidopsis genome or in the genes that dropped out in the first run.

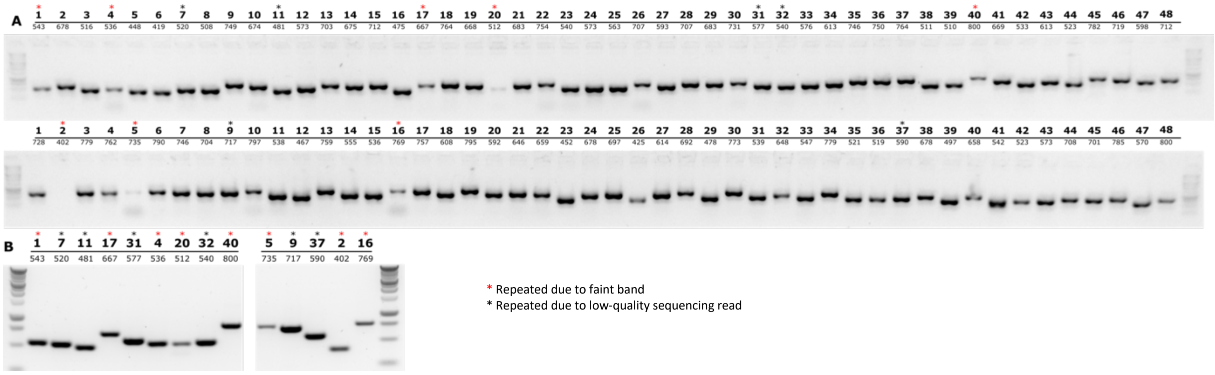


**Figure S5: Amplicon validation by PCR-amplification of a genome-wide Design$_{\text{Sanger,Ext}}$ in Arabidopsis.** To test whether the targets designed for the whole Arabidopsis genome could be amplified and are specific, 48 amplicons were selected from the first run (*SMAP design* run with groups of ± 1000 genes) and 48 amplicons were selected from the dropout-only run (*SMAP design* run with genes that were not retained in the first run but were retained in the second run). The targets were individually PCR amplified, analysed by agarose gel electrophoresis, and Sanger sequenced (**A**). Samples with low intensity bands are indicated with a red asterisk, samples for which a low-quality sequencing read was obtained are indicated with a black asterisk. (**B**) Samples indicated with an asterisk in (A) were amplified again.

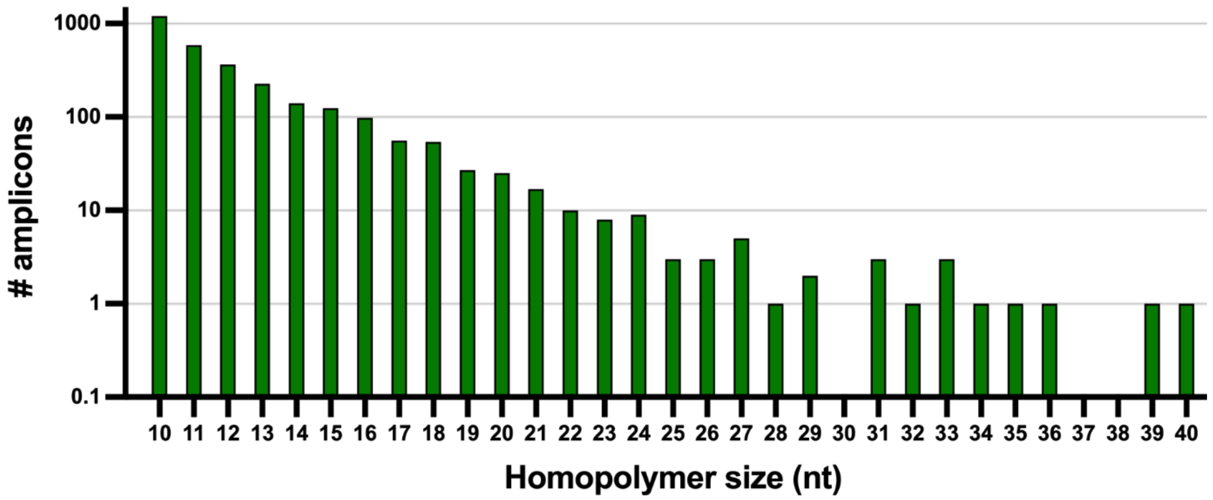


**Figure S6: Number of amplicons designed on the whole genome of Arabidopsis with $Design_{Sanger,Ext}$ with a homopolymer of 10 or more nucleotides.** For each amplicon, the presence of a homopolymer (a stretch of a repeated nucleotide) was counted. Amplicons with more than one homopolymer were counted as 1.

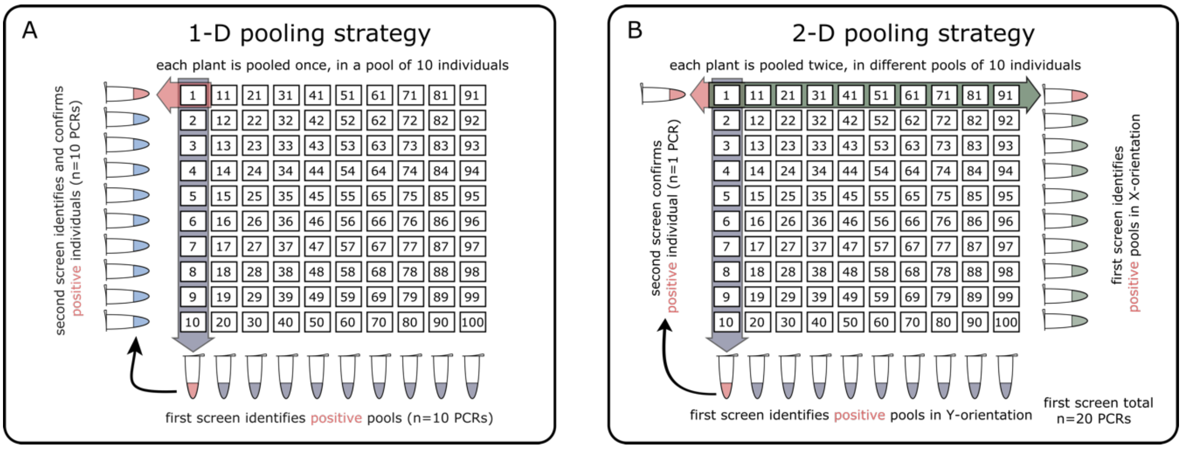


**Figure S7: 1-D and 2-D pooling strategies for genotyping.** Two pooling strategies can be used to decrease the number of PCRs that need to be performed to identify interesting alleles in a population. In a 1-D pooling strategy, each sample is added to one of the pools. Each pool is sequenced in a first screen and the samples of the pools with an allele of interest (here 'positive' pool) are sequenced individually to identify the positive individual. In a 2-D pooling strategy, each sample is added to two of the pools and all pools are sequenced. Positive individuals can be determined through the intersection of the positive pools. A second screen is performed to confirm the positive individual.
